# Interaction of the wireworm species *Agriotes obscurus*, *A. sputator* and *A. lineatus* with a new granule formulation of *Metarhizium brunneum*

**DOI:** 10.1101/2024.12.27.630522

**Authors:** Maximilian Paluch, Tanja Seib, Dietrich Stephan, Andreas Jürgens, Jörn Lehmhus

**Affiliations:** Julius Kühn-Institute (JKI), Federal Research Centre for Cultivated Plants, Institute for Plant Protection in Field Crops and Grassland, Braunschweig, Germany; Julius Kühn-Institute (JKI), Federal Research Centre for Cultivated Plants, Institute for Biological Control, Dossenheim, Germany; Technical University Darmstadt, Department Biology, Chemical Plant Ecology, Darmstadt, Germany

## Abstract

The attract-and-kill strategy is a promising approach in integrated pest management to increase the efficiency of plant protection products by pushing the required contact between target pest and active ingredient. Within the project AgriMet, a new soil granule of the entomopathogenic fungus *Metarhizium brunneum* was developed to control wireworms of the genus *Agriotes* (Coleoptera: Elateridae) in potato cultivation. The formulation is based on autoclaved millet as a substrate to lure the larvae to the infectious conidia on its surface. However, no data existed about the interactions of wireworms with the AgriMet-Granule. Here, I examined the behavioural response of the wireworm species *A. obscurus*, *A. sputator* and *A. lineatus* to the AgriMet-Granule using an olfactometric bioassay in the laboratory. The foraging behaviour of *A. obscurus* and *A. sputator,* with regard to the AgriMet-Granule, was tested with a feed-choice-experiment using a horizontal arena set-up. The tested wireworms significantly preferred autoclaved millet compared to the control. Fungal colonisation on the surface of the AgriMet-Granule did not influence their preference in general. However, differences were observed depending on the respective wireworm species and the two *Metarhizium* spp. isolates used. The feeding bioassay showed that the acceptance of wireworms for the AgriMet-Granule, with a low conidia concentration on its surface, was high. A high conidia concentration on the surface of the AgriMet-Granule led to a decreased acceptance that differed between the wireworm species. However, the survival analysis of wireworms isolated after the feed choice experiment indicated that the wireworms came in contact with the infectious conidia. Overall, the data suggests that autoclaved millet is a suitable substrate for the AgriMet-Granule. It attracts larvae of the genus *Agriotes* to the biocontrol agent using the formulated *M. brunneum* isolate.

## Introduction

Volatile semiochemicals are organic metabolites and play an important role for ecological interactions of very different types of organisms at several trophic levels. Semiochemicals may convey intraspecific (pheromones) or interspecific (allelochemicals) information between microbials, plants and animals, modifying the behaviour of the recipient (Nordlund & Lewis 1976, Agelopoulos et al. 1999, Wyatt 2014). Insects use semiochemicals at all stages of development. For instance, to locate nutrient sources, to find sexual partners (Johnson & Gregory 2006) or to avoid environmental hazards caused by natural antagonists (Davis et al. 2013). In agricultural systems, the release of plant semiochemicals mediates attractive or repellent information for host plant identification by pest insects (Bruce et al. 2005). While attracting semiochemicals may promote the damage by phytophagous insects (Visser 1986, Johnson & Gregory 2006), repellent semiochemicals have been shown to interrupt the identification of their host plants (Hayes et al. 1994). Investigations on the attractive or repellent effects of semiochemicals to pest insects have provided crucial knowledge for effective control strategies in integrated pest management (Noldus 1989). One basic control concept is the behavioural manipulation of target pests using an artificial stimulus to lure it to an attractive source (Cook et al. 2007, Witzgall et al. 2010, Smart et al. 2014). For example, synthetic sex pheromones are used in selective traps to detect and monitor populations of click beetles for risk assessment (Tóth 2013). However, attractants can also be used in combination with plant protection products for pest control. The technique of combining an attractive source with a killing agent is known as “attract and kill” and has been regarded a promising approach for the integrated pest management (El-Sayed et al. 2009).

The “attract and kill” technique has the potential to increase the efficiency of plant protection products by pushing the required contact between active ingredient and target pest (Schumann et al. 2013). In particular, the application of plant protection products against soil-dwelling pests may benefit, as targeted control of phytophagous larvae hidden in the soil is very challenging (Hossain et al. 2007). Soil-dwelling pests in numerous agricultural crops are wireworms, the larval stage of click beetles (Coleoptera: Elaterida) (Parker & Howard 2001, Vernon & Van Herk 2013b). The genus *Agriotes* is the most widespread in Germany with regard to potato cultivation and their typical feeding damage reduces potato quality (Ritter & Richter 2013). Klingler (1957) postulated that the general orientation of wireworm towards the host plant is based on carbon dioxide (CO_2_). In this context, Brandl et al. (2017) evaluated an “attract and kill” strategy against wireworms using CO_2_ emitting capsules. The capsule formulation contained the entomopathogenic fungi *Metarhizium brunneum* (Petch) as biocontrol agent. Brandl et al. (2017) tested the formulation in field trials with varying effectiveness depending on the application technique used. Based on the study of Rastogi et al. (2002), the effectiveness of the capsule formulation could be influenced by microbial, root and faunal respiration of CO_2_ masking the gradient by the capsules to attract wireworms. Nevertheless, biological control of wireworms using *M. brunneum* seems promising and behavioural manipulations through attraction might be beneficial (La Forgia & Verheggen 2019). Since cereal-baited traps were found to be effective for wireworm monitoring (Parker 1996), raw baits should be considered for an “attract and kill” strategy against them.

For this purpose, a new granule formulation of *M. brunneum* was developed to control wireworms as pests in potato cultivation. The idea is that an autoclaved millet grain is coated with liquid fermented biomass of *M. brunneum* by fluidised-bed drying (Bernhardt et al. 2019). The use of millet is justified by its optimal size and stability for common application technology in potato cultivation. The biological activity of the granule should be provided by growth and sporulation out of the thin fungal layer on the surface of the millet grain after application in the soil (Stephan et al. 2020). There, contact of wireworms with the conidia should initialise the infection cycle of *M. brunneum* (Ortiz-Urquiza & Keyhani 2013), which may ultimately result in wireworm reduction (Shah & Pell 2003). Since *Agriotes* larvae are predominately herbivorous (Traugott et al. 2008), an attraction of wireworms by semiochemicals of the substrate millet is conceivable and could increase the likelihood of contact with the infectious conidia on the surface of the granule (El-Sayed et al. 2009). However, no information about an acceptance of wireworms for millet exists. Furthermore, it is unknown if wireworms can olfactorily perceive the natural antagonist *Metarhizium* spp. due to emission of typical semiochemicals. A repellent effect of *Metarhizium* against wireworms would greatly reduce the efficiency of the coated millet control strategy, because the larvae would avoid contact with the active ingredient.

In this study, the interaction of the wireworm species *Agriotes obscurus*, *A. sputator* and *A. lineatus* with a new granule formulation of the entomopathogenic fungus *M. brunneum* was investigated. First, the behavioural response of the mentioned wireworm species to autoclaved millet was assessed using a Y-Olfactometer. Additionally, the fungal coating on the autoclaved millet were taken into account comparing two different isolates (*M. brunneum* isolate JKI-BI-1450, *M. robertsii* isolate JKI-BI-1441) to exclude a repellent effect of the used fungi. Second, the wireworms’ urge to feed on the coated millet grains was examined after different incubation times. Last, the lethal potential of the granule was determined by assessing the mortality of the tested larvae after the feed choice experiment. This investigation aims to fill the gap on the behavioural response and the feeding behaviour of different wireworm species to a millet-based mycoinsecticide, which is crucial for the optimization of future control strategies against them.

## Material and Methods

### Wireworm breeding for the bioassays

Larvae of the species *Agriotes obscurus*, *A. sputator* and *A. lineatus* emerged from the laboratory breeding at the JKI Institute for Plant Protection in Field Crops and Grassland (Braunschweig, Germany). Click beetles of the respective species were collected in Wohld (52°18’11.0”N 10°41’11.6”E, Germany) and determined to species level using the identification key of Lohse (1979). Afterwards, beetles were transferred to buckets with soil and wheat (*Triticum aestivum*, Cultivar: Primus, Deutsche Saatgutveredelung AG, Lippstadt, Germany) at 20 °C for laying eggs following the protocol of Kölliker et al. (2009) with minor modifications. After about eight months, larvae originated from the breeding were removed and stored in plastic boxes (18.3 x 13.6 x 6.4 cm, Baumann Saatzuchtbedarf, Waldenburg, Germany) with a moist paper towel at 5 °C until use. To ensure healthy and vital wireworms, the larvae were transferred into soil at 15 °C for acclimatisation and fed with *Triticum aestivum* seeds (Cultivar: Primus, Deutsche Saatgutveredelung AG, Lippstadt, Germany) fourteen days before the start of the experiment.

### AgriMet-Granule

The AgriMet-Granule is an autoclaved millet grain coated with the *Metarhizium brunneum* isolate JKI-BI-1450, isolated in 2016 from an infected *A. lineatus* beetle in Germany. For mass production of the fungus material, a liquid fermentation was conducted and the biomass including submerged spores, hyphae and secondary metabolites was coated with the help of a fluidised bed dryer on autoclaved millet grains of the species *Setaria italica* (Alnatura Produktions-und Handels GmbH, Darmstadt, Germany). To investigate the influence of the formulated *Metarhizium* isolate, a comparable granule was produced using the *M. robertsii* isolate JKI-BI-1441 originated from an *Agriotes* sp. larva gathered during field surveys in Italy. Both granules were provided by Tanja Bernhardt (JKI Institute for Biological Control, Darmstadt, Germany) and completely dry, while 1 kg contained 4.5 g of dry fungal biomass. To ensure the functionality, storage at 5 °C and darkness was no longer than twenty weeks.

### Olfactometric bioassay with the AgriMet-Granule

#### Y-glass tube for the olfactometric bioassay

The olfactory perception of the *Metarhizium brunneum* isolate JKI-BI-1450 and *M. robertsii* isolate JKI-BI-1441 grown on millet by the wireworm species *A. obscurus*, *A. sputator* and *A. lineatus* were examined using a Y-glass tube (W.O. Schmidt GmbH Laboratoriumsbedarf, Braunschweig, Germany) shown in Figure 3. The total length of the tube was 150 mm with an internal diameter of 35 mm. Parafilm was placed on each of the three openings to prevent the escape of the wireworms and the volatile organic compounds. The glass tube was filled with vermiculite (Floragard, Oldenburg, Germany) with a grain size of 2-3 mm. In order to obtain a homogenous substrate and to avoid clumping, the vermiculite was additionally sieved (1 mm mesh) before use. After that, 40 g vermiculite were moistened with 25 ml tap water for each glass tube to ensure adequate conditions for wireworm movement.

**Figure 1:**
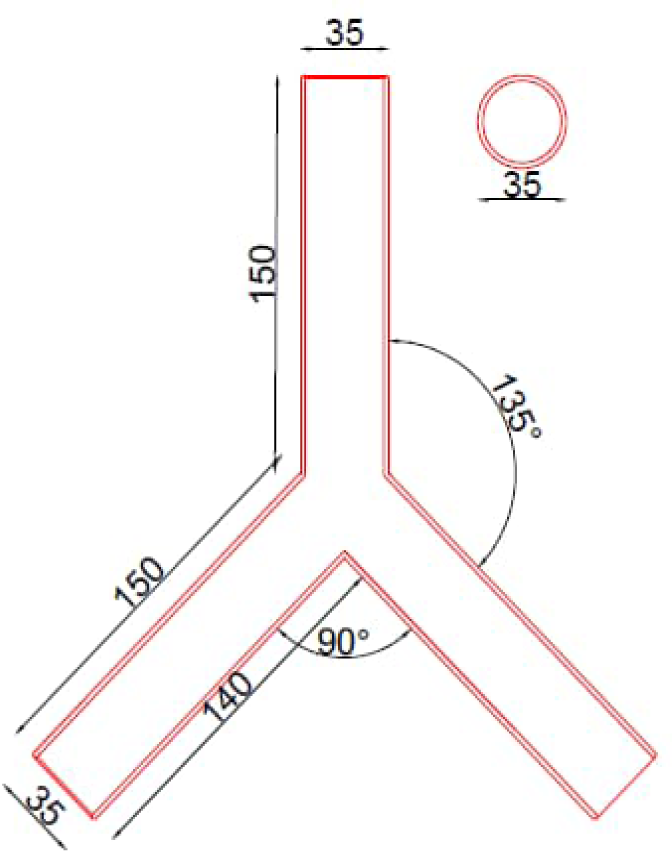
Schematic illustration of the Y-glass tube for the olfactrometric bioassay about the behavioural response of wireworms to the AgriMet-Granule.

**Figure 2:**
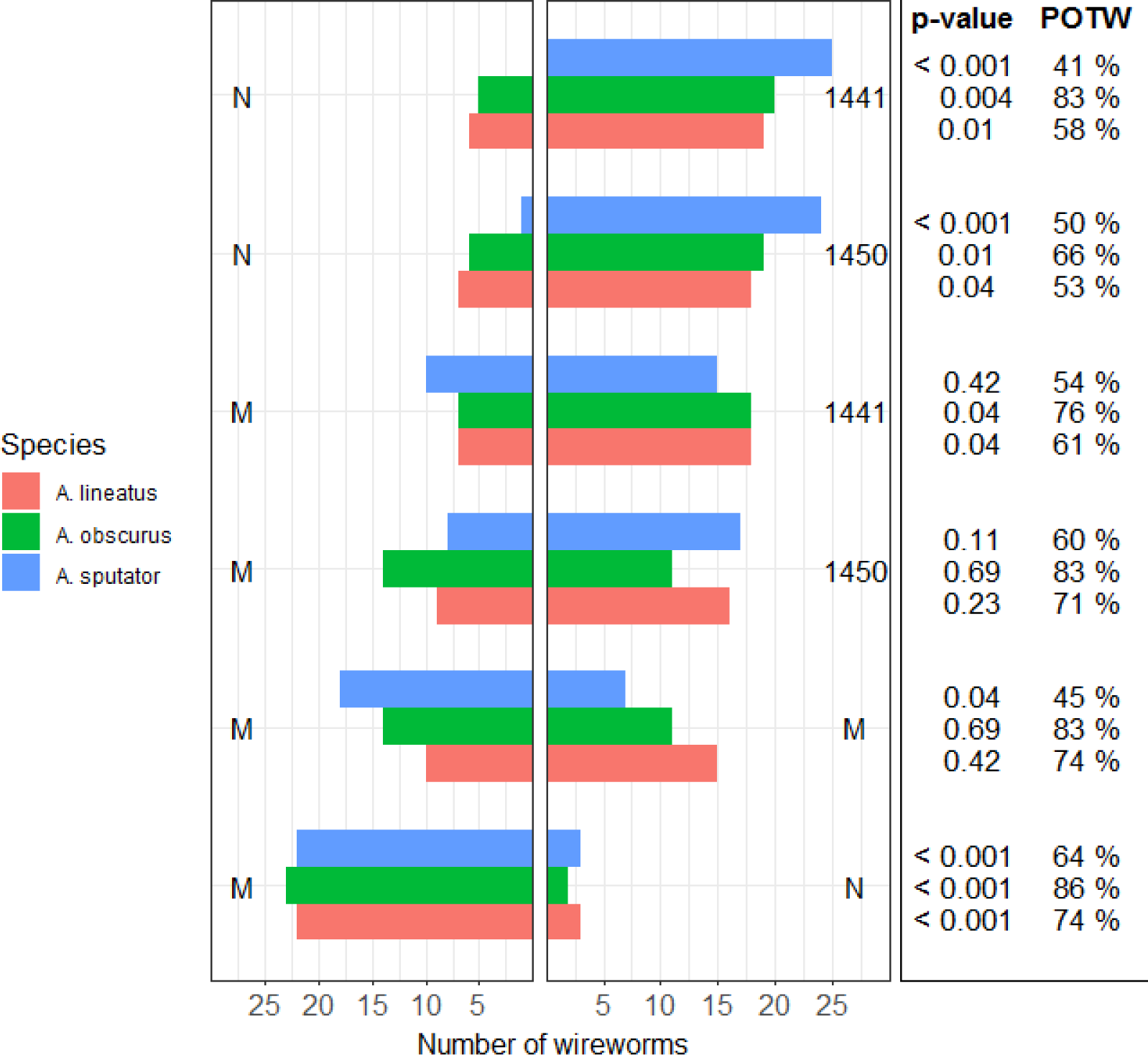
Number of wireworms reaching one side of the Y-glass tube after 2 h. The Y-glass tube was filled with different offers at the two ends (M = autoclaved millet, 1450 = *M. brunneum* isolate JKI-BI-1450 grown on autoclaved millet, 1441 = *M. robertsii* isolate JKI-BI-1441 grown on autoclaved millet, N = nothing). The colours of bars indicate the wireworm species *Agriotes lineatus* (red), *A. obscurus* (green) and *A. sputator* (blue). The *p*-values are based on a two-tailed Exact Binomial test (*α* = 0.05) and are shown on the right as well as the proportion of all tested wireworms who moved and made a decision (POTW).

**Figure 3:**
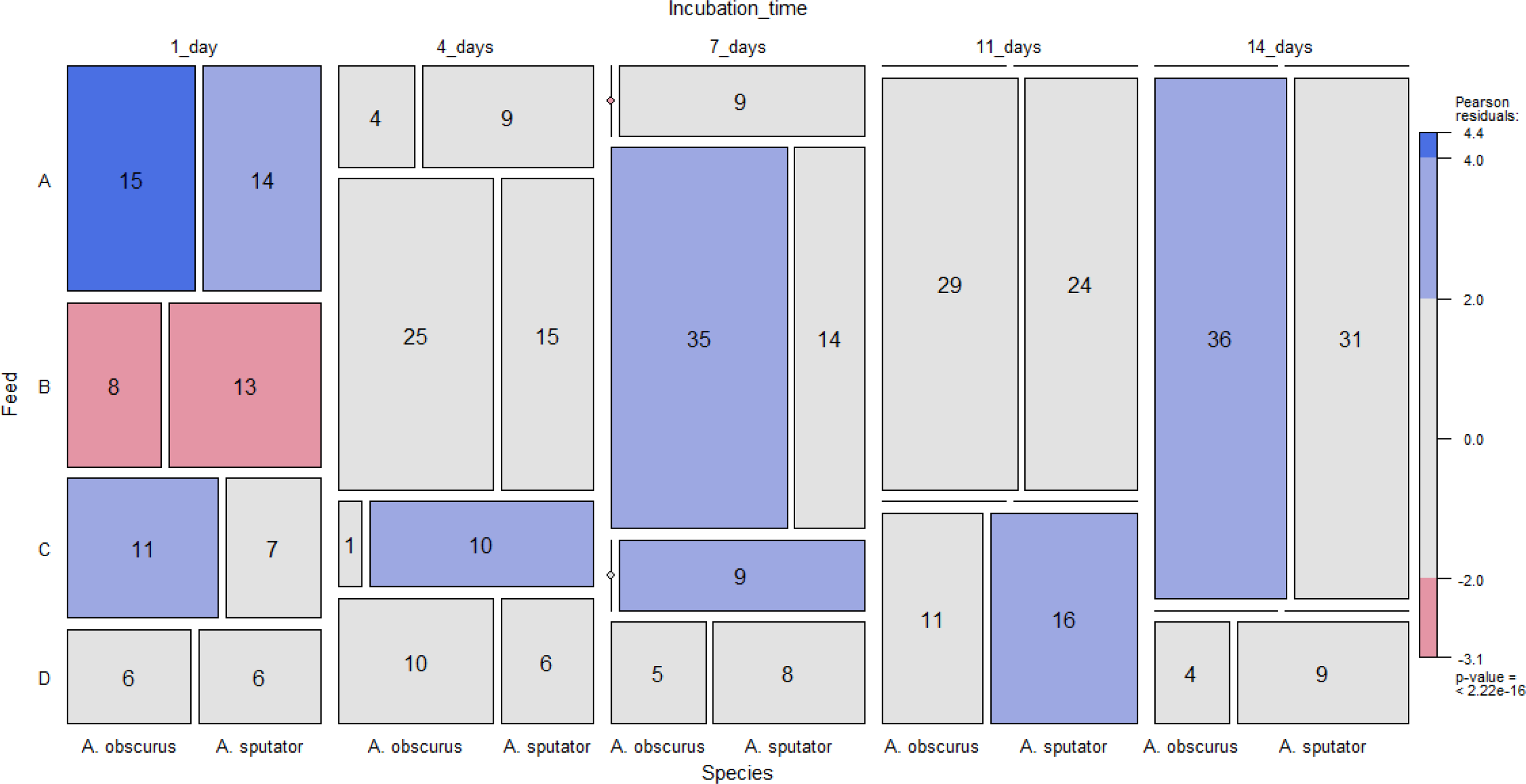
Three-way mosaic plot reflecting the presence of feeding marks (A: AgriMet-Granule, B: wheat, C: AgriMet-Granule and wheat, D: none) of the tested wireworm species (*A. obscurus*, *A. sputator*) based on different incubation times of the AgriMet-Granule (1_day, 4_days, 7_days, 11_days, 14_days). Height and width of the boxes correspond to the frequency of feeding marks of the respective wireworm species in each variant. The colours correspond to the size of the Pearson residuals. High deviations are presented deep blue or red (> 4, corresponds to *α* = 0.0001), medium deviations light blue or red (< 4 and > 2, corresponds to *α* = 0.05) and smal deviations grey (< 2).

#### Experimental set-up of the olfactometric bioassay

The dual-choice bioassay consisted of the following treatments: (A) autoclaved millet – autoclaved millet, (B) JKI-BI-1450 – autoclaved millet, (C) JKI-BI-1450 – nothing, (D) JKI-BI-1441 – autoclaved millet, (E) JKI-BI-1441 – nothing. Treatments were carried out simultaneously with the wireworm species *A. obscurus*, *A. sputator and A. lineatus* which led to a total of fifteen glass tubes. Each glass tube was filled with the AgriMet-Granule of the corresponding fungal isolate, autoclaved millet or nothing opposite against each other in the two openings at the top. Prior to use, the AgriMet-Granule was incubated for 14 days at 25 °C on 1.5 % water agar to ensure fungal colonisation on the surface. The conidia concentration was determined as in the experimental set-up of the feeding bioassay and reached 8.41x10^7^ conidia grain^-1^ (±SD 5.43x10^6^) for Isolate JKI-BI-1450 and 2.35x10^8^ conidia grain^-1^ (±SD 9.23x10^6^) for isolate JKI-BI-1441. Aliquots weighing 1 g of the respective AgriMet-Granule or autoclaved millet were prepared and moistened with 500 µl autoclaved tap water 24 h before the experiment in order to simulate the swelling in the soil. Wireworms were inserted at the opening at the bottom 20 min after bait insertion to ensure the distribution of volatile organic compounds. After 2 h at 20 °C in the dark, the location of wireworms was recorded. Only those larvae that reached at least the bottom end of one of the two sides connected to the openings at the top were used for the evaluation. No airflow was installed and the experiment was repeated until twenty-five larvae of each treatment had moved and made a decision.

### Feeding bioassay with the AgriMet-Granule

#### Trial arena of the feeding bioassay

The urge of the wireworm species *A. obscurus* and *A. sputator* to feed on millet grains coated with the fungal isolate JKI-BI-1450 was examined by a choice experiment. For this purpose, Petri dish lids of two different manufacturers (Greiner Bio-One GmbH, Kremsmünster, Austria; Brand GmbH & Co. KG, Wertheim, Germany) were used in order to create a completely closed arena with a diameter of 15 cm and a height of 1 cm. The low height was necessary to be able to observe the movement of the wireworms during trials. As substrate 1 kg of a 5:1 mixture of potting soil (Einheitserde Classic Pikiererde CL P, Gebrüder Patzer GmbH & Co. KG, Sinntal, Germany) and sand moistened with 300 ml tap water (49.2 % residual moisture) was used.

#### Experimental set-up of the feeding bioassay

The feed choice experiment was carried out placing the same amount (g) of the AgriMet-Granule and *Triticum aestivum* seeds (Cultivar: Primus, Deutsche Saatgutveredelung AG, Lippstadt, Germany) opposite against each other in the trial arena and releasing one wireworm in the middle for 48 h. After 24, 25, 26, 27, 28, 29 and 48 h the location of the wireworm was observed and assessed as follows: (W) wireworm in contact with wheat, (G) wireworm in contact with the AgriMet-Granule, (N) wireworm in no contact with wheat or the AgriMet-Granule. At the end, the presence of feeding marks on the AgriMet-Granule (A), wheat (B), both (C) or none (D) was examined with the help of a binocular (Carl Zeiss AG, Oberkochen, Germany). Wheat was used because wireworms accept it very well, while potatoes were not suitable for the test system due to the large size. The AgriMet-Granule was offered after five different incubation times to investigate the influence of the fungal colonisation on the surface of the millet. The coated millet was incubated for 1, 4, 7, 11 or 14 days at 25 °C on 1.5 % water agar and fungal growth was defined by the number of conidia grain^-1^ millet. Therefore, ten millet grains of the respective incubation time were washed with 0.1 % Tween80® and conidia were removed by vortexing to determine the concentration of conidia grain^-1^ with a haemocytometer. The examination of the conidia concentration was repeated five times. Twenty trial arenas were used at the same time and stored at 20 °C in darkness. At the end of the experiment, tested wireworms were isolated in small cans with moist paper towel and incubated at 25 °C and darkness for six weeks to assess a possible infection with the *M. brunneum* isolate JKI-BI-1450. The larvae were feed with wheat (Cultivar: Primus, Deutsche Saatgutveredelung AG, Lippstadt, Germany) to avoid starving. Infected wireworms were identified based on fungal outgrowth from the cadaver and on morphological criteria of *Metarhizium* spp. (Zimmermann 2007).

### Statistical analyses

Statistical analyses were done using the software R Studio (Version 1.4.1106) (RStudio Team 2020). Selection of the best-fitted model was based on the Akaike Information Criterion (AIC) (Burnham & Anderson 2002) after backward elimination of the full model. Subsequent model diagnostic was carried out by visual inspection of the QQ-Plot (sample quantile-theoretical quantile) and Residuals-Prediction-Plot to confirm normal distribution and variance homogeneity.

The influence of the incubation time (explanatory variable) on the conidia concentration on the surface of the AgriMet-Granule (target variable) was analysed with a Linear Model (LM). After log transformation of the target variable, normal distribution and variance homogeneity were met. If significant effects were identified in the analysis of variance (ANOVA, *α* = 0.05), differences between the respective incubation times were examined using the post hoc Tukey HSD test (*α* = 0.05) included in the *emmeans* R package (Lenth et al. 2018).

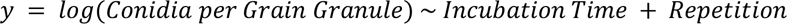

To analyse the preference of the tested wireworm species for autoclaved millet or the AgriMet-Granule, a two-tailed Exact Binomial test (*α* = 0.05) was performed for each comparison individually.

The feeding activity of the tested wireworm species was visualised with a three-way mosaic plot using the *vcd* R package (Meyer et al. 2020). Height and width of the boxes correspond to the frequency of feeding marks of the respective wireworm species in each variant. The colours shown in the mosaic plot refer to the Pearson residuals and thus reflect patterns of dependency. Blue means that there are more observations in the box than would be expected under the null model, and red means that there are fewer observations than would have been expected. The intensity of the colour corresponds to the size of the Pearson residuals.

The visits of the tested wireworm species to the AgriMet-Granule or wheat were expressed as proportion of the seven possible observation time points. The influence of the incubation time of the AgriMet-Granule (explanatory variable) on the visits (target variable) was analysed by using a Generalised Linear Model (GLM, binominal distribution). Global effects were determined with an analysis of deviance. The pairwise comparisons were conducted as described above.

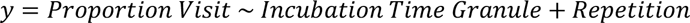

Kaplan-Meier-Analysis was used to determine the survival probability of the tested *A. obscurus* and *A. sputator* larvae over time subsequent to the feed-choice-experiment. Survival curves over time were created using the *survival* R package (Therneau 2021) and compared with the log-rank test for global differences. To detect significant differences between the five variants within a wireworm species a pairwise comparison of survival curves based on the Bonferroni method was carried out using the *survminer* R package (Kassambara et al. 2021). Mycosis of wireworms was recorded as an “event”. If no event was observed by the end of the study, the total survival time could not be accurately determined and was censored.

All graphs except the mosaic plot were created with the R packages *ggplot2* (Wickham 2016), *ggpubr* (Kassambara 2020), *RColorBrewer* (Neuwirth 2014), and *multcompView* (Graves et al. 2019).

## Results

### Conidia per grain AgriMet-Granule

The conidia concentration on the surface of the AgriMet-Granule coated with *Metarhizium brunneum* isolate JKI-BI-1450 was determined by the number of conidia grain^-1^ AgriMet-Granule and is presented in Table 1. After one day of incubation, no fungal outgrowth was observed and 5.89x10^3^ conidia grain^-1^ (±SD 5.06x10^2^ conidia grain^-1^) were detached. Three days later, the number of conidia grain^-1^ increased only slightly to 8.11x10^3^ (±SD 6.98x10^2^ conidia grain^-1^), but *M. brunneum* started to grow and very fine hyphae could be observed. After seven days of incubation, the conidia concentration increased significant (Tukey HSD test, *p* < 0.0001) to 1.96x10^7^ conidia grain^-1^ (±SD 1.68x10^6^ conidia grain^-1^). The fungal growth of *M. brunneum* on the surface of the AgriMet-Granule was clearly visible based to the white mycelium and the green conidia. The fungus continued to significantly grow (7-11 days: Tukey HSD test, *p* < 0.0001) on the surface of the AgriMet-Granule and reached and maximum of 7.42x10^7^ conidia grain^-1^ (±SD 6.39x10^6^ conidia grain^-1^) after 14 days. At this point, the grain was completely covered with conidia of *M. brunneum*.

**Table 1:** Adjusted mean values of *M. brunneum* conida per grain AgriMet-Granule and standard deviation (±SD) after 1, 4, 7, 11 and 14 days of incubation on 1.5 % water agar at 25 °C and darkness. Adjusted mean values with the same letters are not significantly different based on LM (y = log(Conidia per Grain Granule) ∼ Incubation Time + Repetition) and Tukey HSD test (*α* = 0.05).

| Incubation time [days] | Number of conidia grain<sup>-1</sup> AgriMet-Granule |  |  |
| --- | --- | --- | --- |
| | adjusted mean | ±SD | sig |
| 1 | 5.89x10<sup>3</sup> | 5.06x10<sup>2</sup> | a |
| 4 | 8.11x10<sup>3</sup> | 6.98x10<sup>2</sup> | a |
| 7 | 1.96x10<sup>7</sup> | 1.68x10<sup>6</sup> | b |
| 11 | 5.16x10<sup>7</sup> | 4.44x10<sup>6</sup> | c |
| 14 | 7.42x10<sup>7</sup> | 6.39x10<sup>6</sup> | c |

### Olfactometric bioassay with the AgriMet-Granules

The behavioural response to autoclaved millet and/or the AgriMet-Granule examined with a Y-glass tube did not differ between *Agriotes obscurus*, *A. sputator* and *A. lineatus*. Figure 4 illustrates that neither the *Metarhizium brunneum* isolate JKI-BI-1450 nor the *M. robertsii* isolate JKI-BI-1441 had a repellent effect on the tested wireworm species. On the contrary, *A. obscurus* and *A. lineatus* were significantly attracted to the isolate JKI-BI-1441 compared to autoclaved millet (Exact-Binomial-Test, *p* = 0.04). In addition, both isolates were significantly preferred by all wireworm species when the isolates were offered in comparison to nothing (Exact-Binomial-Test, *p* < 0.05). The simultaneous offer of the isolate JKI-BI-1450 and autoclaved millet resulted in a more or less even split between the twenty-five wireworms of the respective species tested (Exact-Binomial-Test, *p* > 0.1). Similar results were observed with autoclaved millet at both ends of the Y-glass tube. All tested wireworm species were significantly attracted to autoclaved millet compared to nothing (Exact-Binomial-Test, *p* < 0.001). Although the behavioural response was comparable between the species, they differed in their activity. The proportion of all tested wireworms who moved and made a decision was highest by *A. obscurus* in each comparison and ranged between 66-86 %. Larvae of *A. sputator* showed the lowest activity with a maximum of 60 %. The comparison between the isolate JKI-BI-1441 and nothing with *A. sputator* resulted in the lowest proportion of 41 %.

**Figure 4:**
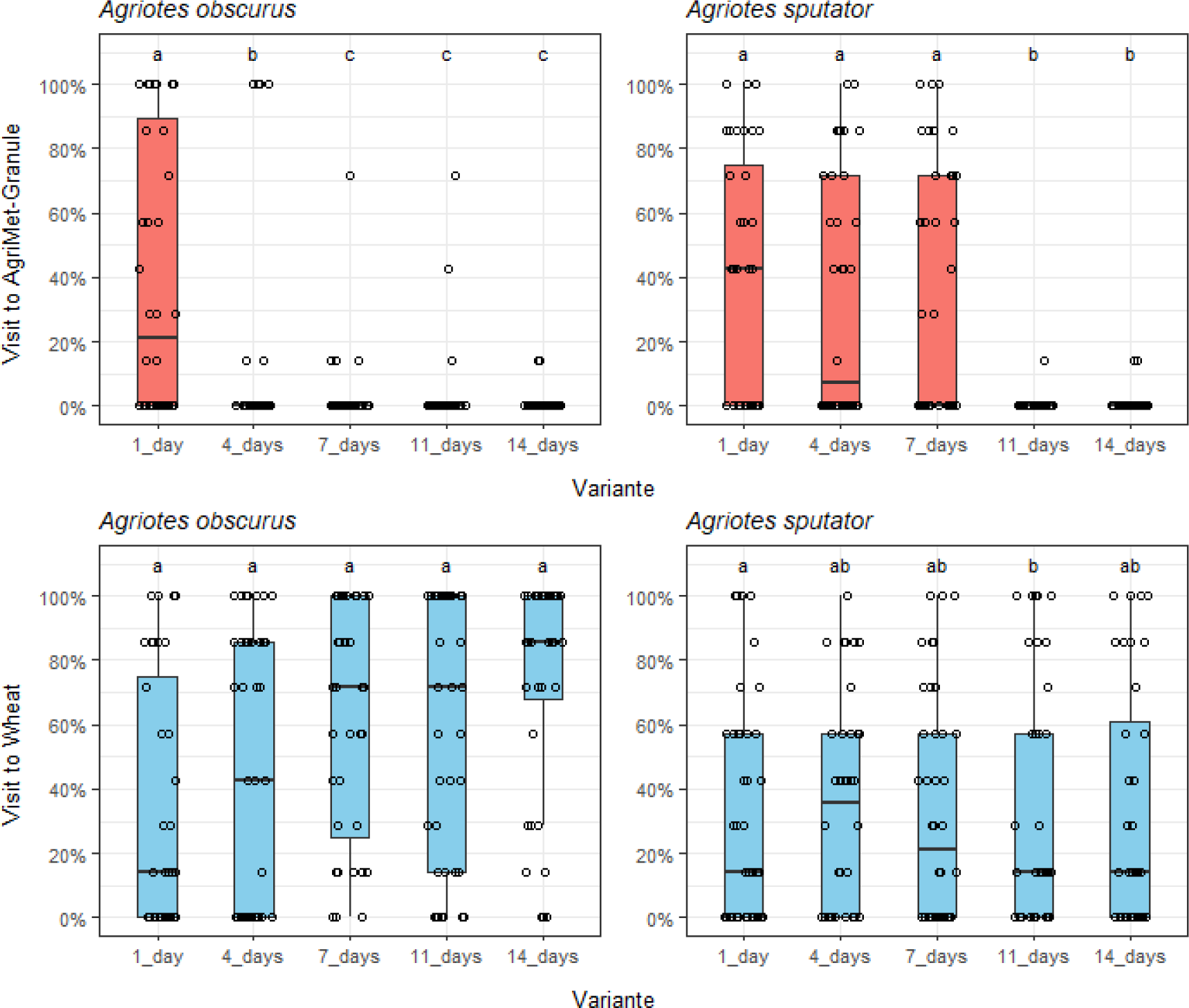
Boxplots of the visits (proportion of 7 observations in percent) to the AgriMet-Granule in red or wheat in blue of the wireworm species *Agriotes obscurus* and *A. sputator* in five different variants (1_day = 1 day incubated AgriMet-Granule against wheat, 4_days = 4 days incubated AgriMet-Granule against wheat, 7_days = 7 days incubated AgriMet-Granule against wheat, 11_days = 11 days incubated AgriMet-Granule against wheat, 14_days = 14 days incubated AgriMet-Granule against wheat). The boxes illustrate the median, 25 % and 75 % quantile. Jittered points represent each larva. Variants with the same letters within the respective species and food source are not significantly different. GLM (y = Proportion Visit ∼ Incubation Time Granule + Repetition, family=binomial), Tukey HSD test (*α* = 0.05).

### Feeding bioassay with the AgriMet-Granule

A feed choice experiment was carried out to investigate the urge of *Agriotes obscurus* and *A. sputator* to feed on the AgriMet-Granule coated with *Metarhizium brunneum* isolate JKI-BI-1450. The results of the feeding assay are shown in a mosaic plot (Figure 5). The AgriMet-Granule was well accpeted by both wireworm species after one day of incubation and the frequency of feeding marks on the AgriMet-Granule (A and C) was higher than expected. Compared to the other variants, wheat (B) was the least prefered in the 1 day variant. After 4 days of incubation of the AgriMet-Granule, the frequency of feeding marks on the AgriMet-Granule by *A. obscurus* decreased from 26 to 5 larvae, wherase the amount of *A. sputator* larvae remained constant at 19. In addition, the acceptance of wheat increased due to the raising amount of feeding marks. After 7 days of incubation, the AgriMet-Granule was no longer accpeted as food source by *A. obscurus* and the frequency of feeding marks on wheat was higher than expected with 35 larvae. However, *A. sputator* was still feeding on the AgriMet-Granule (18 larvae) and the frequency of feeding marks on wheat was smaler compared to *A. obscurus*. After an incubation time of 11 days, no feeding marks on the AgriMet-Granule could be assessed by both wireworm species. The frequency of no feeding marks on both offers (D) increased and was higher than expected with 16 larvae by *A. sputator*. After 14 days of incubation, wheat was the only food source that was accepted by both wireworm species, but the share of no feeding marks decreased again.

**Figure 5:**
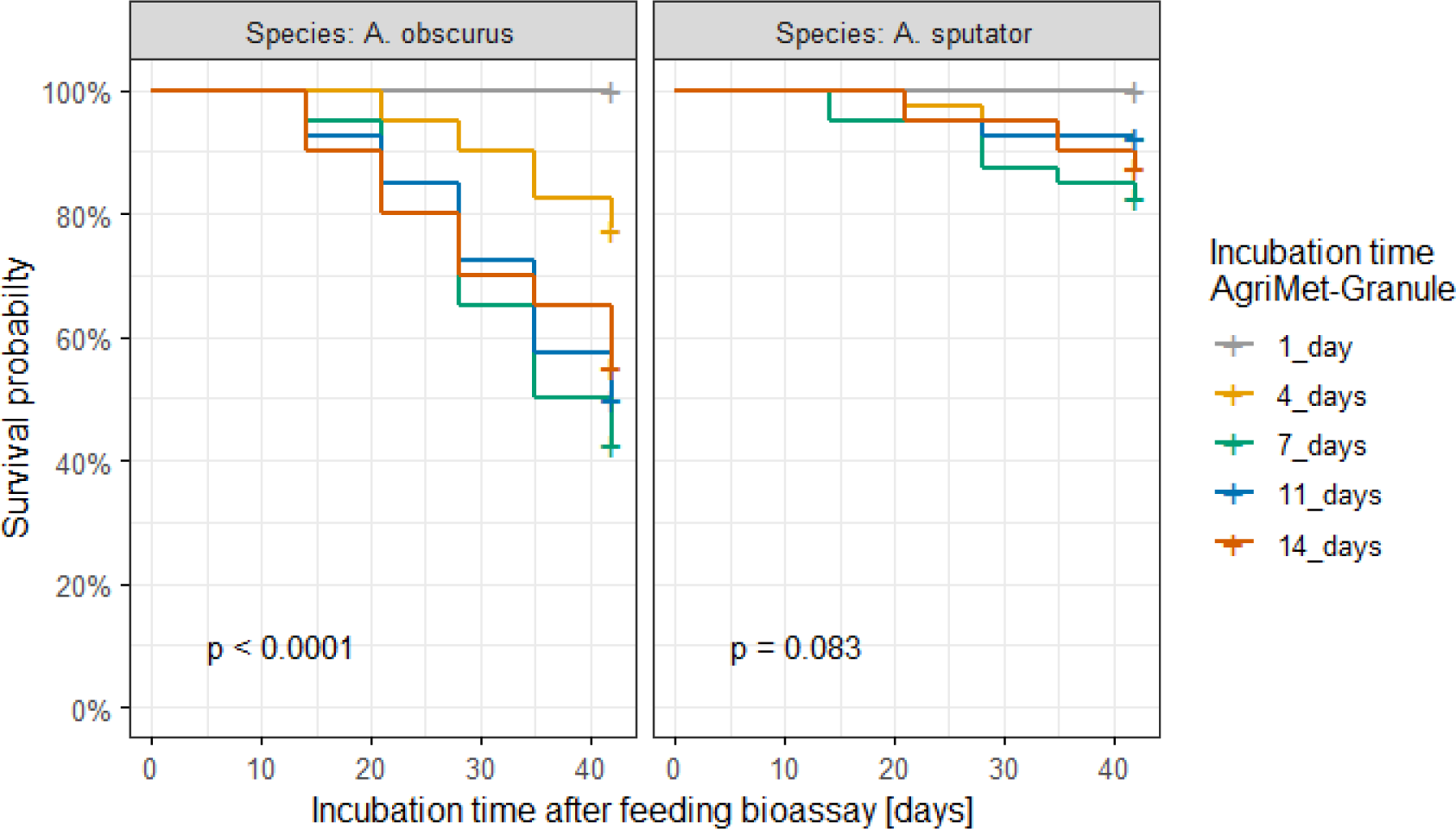
Overall survival probability in percent (Kaplan-Meier-Analysis) of the wireworm species *Agriotes obscurus* and *A. sputator* isolated after the feed-choice bioassay with wheat and the AgriMet-Granule with different incubation times and conidia concentrations (1_day = 5.89x10^3^ conidia grain^-1^; 4_days = 8.11x10^3^ conidia grain^-1^; 7_days = 1.96x10^7^ conidia grain^-1^; 11_days = 5.16x10^7^ conidia grain^-1^; 14_days = 7.42x10^7^ conidia grain^-1^). Larvae were incubated individually for 42 days under laboratory conditions at 25 °C and darkness in small cans filled with moist paper towel. The statistic of the shown *p*-values is based on the log-rank test (*α* = 0.05) and indicates differences between the variants within the respective wireworm species.

The assessment of visits at the AgriMet-Granule or wheat 24, 25, 26, 27, 28, 29 and 48 h after the wireworms were released in the trial area is shown in Figure 6. *Agriotes obscurus* and *A. sputator* moved between the offers and were observed at the AgriMet-Granule and wheat in the 1_day variant. After 4 days of incubation, the visits at the AgriMet-Granule of *A. obscurus* decreased significantly, whereas *A. sputator* was still monitored. With a few exceptions, *A. obscurus* was no longer seen at the AgriMet-Granule incubated for 7, 11 and 14 days. However, *A. sputator* was observed at the AgriMet-Granule incubated for 7 days, but only marginally after 11 or 14 days. The wireworms which could not be observed at the AgriMet-Granule in the variants mentioned were active due to the observation at wheat during all incubation times.

Wireworms that were not seen at the AgriMet-Granule at higher incubation times must have been there due to the overall survival probability shown in Figure 7. The incubation time of the AgriMet-Granule had a significant influence on the survival probability of *A. obscurus* isolated after the feeding assay (log-rank test, *p* < 0.0001). 20 of the 40 tested wireworms exposed to the AgriMet-Granule incubated for 7 days were killed by the *M. brunneum* isolate JKI-BI-1450. The survival probability was under 50 % after the entire observation time of 42 days. The incubation time of 11 and 14 days of the AgriMet-Granule led to 17 and 14 larvae with a mycosis with no significant difference to the 7 days variant (Bonferroni, *p* > 0.05). Four days of incubation of the AgriMet-Granule resulted in a significantly higher survival probability of approximate 75 % compared to seven days of incubation (Bonferroni, *p* = 0.0096). No mycosis was observed at both wireworm species after 1 day of incubation of the AgriMet-Granule. However, the survival probability of *A. sputator* was not influenced by the incubation time of the AgriMet-Granule (log-rank test, *p* = 0.083). Only 3-6 larvae of *A. sputator* were killed by the *M. brunneum* Isolate JKI-BI-1450 42 days after the feeding assay and the survival probability was not under 80 % in any variant.

## Discussion

In the development of an efficient mycoinsecticide, a higher probability of contact between target pest and fungal biocontrol agent can be achieved by the “attract-and-kill” technique (Schumann et al. 2013, Vemmer et al. 2016). Using a bait as substrate for the formulation that is well accepted as nutrient source may serve this purpose. Autoclaved millet was used for the AgriMet-Granule intended for the control of wireworms in potato cultivation. The attractive effect of autoclaved millet was confirmed by a clear preference of the tested wireworm species *A. obscurus*, *A. sputator* and *A. lineatus* in the olfactometric bioassay compared to the control. Semiochemicals emitted by the autoclaved millet grains might be responsible for the preference of the larvae. The perception and orientation of soil-dwelling pests towards a host plant by a CO_2_ gradient is widely reported for belowground arthropods (Johnson & Gregory 2006; Guerenstein & Hildebrand 2008). For example, larvae of the western corn rootworm (*Diabrotica virgifera*) rely mainly on CO_2_ as attractive semiochemical during foraging (Bernklau & Bjostad 1998). Behavioural studies of Doane & Klingler (1978) showed that *A. obscurus* and *A. lineatus* perceive CO_2_. Using electron microscopy, they found that clusters of sensilla on the labial and maxillary palps serve this perception. Since germination and associated mitochondrial metabolism of the millet grains is absent due to autoclaving, the production of CO_2_ was reduced (Bewley & Black 2012). Thus, an important attractive volatile might have been missing. However, the exact role of CO_2_ for arthropods is being discussed and probably only important for the initialisation of foraging behaviour (Johnson & Nielsen 2012). Even without the stimulus of CO_2_, wireworms are able to orient themselves towards specific volatile semiochemicals. Root produced volatile aldehydes including Hexanal and (E)-Hex-2-enal extracted from barley attracted *A. sordidus* in a comparable olfactometric assay to the one used here (Barsics et al. 2017). A gas chromatography–mass spectrometry analysis (GCMS) of milled millet revealed, that the proportion of aldehydes was the highest with 21.81–34.21 % and derivates of Hexanal were the most abundant (Liu et al. 2012). Similar volatile aldehydes were also found in autoclaved seeds (Stotzky & Schenck 1976). Thus, Hexanal derivates are very likely the reason for the preference of wireworms to autoclaved millet in each comparison of the olfactometric bioassay performed in this study.

However, the AgriMet-Granule is coated with a thin layer of the entomopathogenic *Metarhizium brunneum*, which is intended to proliferate under suitable conditions providing lethal potential against wireworms. Therefore, the wireworm acceptance with respect to autoclaved millet colonised with the natural antagonist *Metarhizium* spp. on the surface was investigated in a parallel bioassay. Comparing autoclaved millet with autoclaved millet completely covered with conidia of *M. brunneum* isolate JKI-BI-1450 or *M. robertsii* isolate JKI-BI-1441 clarified that the preference for autoclaved millet was not negatively influenced by fungal growth. These results are in contrast to the those of Mburu et al. (2009), who investigated the relationship between the virulence and repellency of *M. anisopliae* to the termite species *Macrotermes michaelseni* in a glass Y-tube. They reported a dose-related repellency of virulent isolates. In addition, they identified differences in the volatile blends of two *Metarhizium* isolates, which influenced the behavioural response of the tested termites (Mburu et al. 2011). Bojke et al. (2018) also confirmed the unique volatile blends of different *Metarhizium* spp. isolates. The olfactometric bioassay in this study showed an increased preference for the *M. robertsii* isolate JKI-BI-1441 over the *M. brunneum* isolate JKI-BI-1450. The occurrence of unique volatile blends of the isolates might be an explanation for this. The tested wireworms might have perceived the different blends, which resulted in a varied behavioural response towards the isolates. The exact volatile profile of the used isolates is yet unknown. Mburu et al. (2011) identified qualitative and quantitative differences in the volatile profile of two *Metarhizium* spp. isolates investigated. Although Furanone was rated as the most repellent substance against termites, they pointed out the importance of the composition and the relative amounts of substances of volatile blends. Degradation of millet during the assimilation of entomopathogenic fungi could lead to a quantitative increase of volatile aldehydes like Hexanal derivatives, thereby altering the relative amounts of the compounds. Nevertheless, the AgriMet-Granule can attract *A. obscurus*, *A. sputator* and *A. lineatus*. However, attraction is only the first part of the intended “attract-and-kill” strategy to control wireworms using the AgriMet-Granule. Prerequisite for the initialisation of the lethal infection cycle is the direct contact between infectious conidia of *Metarhizium* spp. and the larvae (Goettel & Glare 2010). The urge of wireworms to feed on the AgriMet-Granule can fulfil the required contact and is discussed in the following section.

Rizzo & Lehmhus (2014) demonstrated that wireworms are generalist herbivores. Therefore, the acceptance of autoclaved millet as food source was expected and confirmed in the feeding bioassay with *A. obscurus* and *A. sputator*. The larvae bite into the AgriMet-Granule with almost no fungal growth or with 8.11x10^3^ conidia grain^-1^ on the surface. However, the urge to feed on the AgriMet-Granule was no longer observed at concentrations of 5.16x10^7^ conidia grain^-1^. Again, the olfactory perception of volatile blends could have influenced the behavioural response. Since wireworms can make use of a variety of different nutrient resources, specific volatiles are not required for host identification. Rather, the right composition and ratio of volatiles might be responsible for host recognition (Bruce & Pickett 2011). Thorpe et al. (1947) showed that plant juices or solutions containing carbohydrate, fatty or protein substances elicit the biting response of wireworms. In this context, the initial stimulus to bite into the millet grain might be masked by the volatile blend of the *M. brunneum* isolate used. Consequently, direct contact to the AgriMet-Granule through food ingestion by wireworms might only be ensured until proliferation of *M. brunneum* on the autoclaved millet reaches a certain conidia concentration.

The conidia concentration is of great importance for the intended control strategy, because it is closely linked to the lethal potential of *Metarhizium* spp. (Gabarty et al. 2014). The AgriMet-Granule with a concentration of 1.96x10^7^ conidia grain^-1^ was accepted as food source by *A. obscures* and the survival probability was under 50 %. In contrast, mortality of *A. sputator* was relatively low with a survival probability of 83 %, indicating a selective effectiveness of *M. brunneum* isolate JKI-BI-1450 against different wireworm species. Surprisingly, the survival probability of *A. obscurus* also decreased significantly in the bioassay where no feeding scars on the AgriMet-Granule were observed. Larval mortality is based on contact with the AgriMet-Granule. Although no feeding scars were found, contact alone, and not necessarily feeding, appears to be sufficient for the desired lethal effect. Overall, under laboratory conditions, the AgriMet-Granule showed potential for wireworm control through its attracting effect, acceptance as nutrient source, and formation of a lethal conidia concentration of *M. brunneum* on the surface.

The results suggest the control potential of wireworms by the AgriMet-Granule based on laboratory set ups, but no further environmentally influencing factors were taken into account. However, potatoes are grown in different regions with different soil types, so the emission of further interfering semiochemicals is to be expected. The diffusivity of volatile semiochemicals in terms of distance and intensity depends on the physical properties of soil (Rolston & Møldrup 2012), which get influenced by the proportion of silt, clay, sand and organic matter (Aochi & Farmer 2005). Sand and organic matter increase the formation of soil pores for an optimised diffusion of gases, whereas a high proportion of clay and loam compacts the soil (Boyle et al. 1989). The soil structure of a potato ridge is related to the region and sandy soils are preferred in Germany because of the capacity to warm up well and their loose structure (Paffrath et al. 2004). In this context, the volatile semiochemicals of the AgriMet-Granule may attract wireworms over a greater distance, which should be investigated in further experiments. However, there are ubiquitous bacteria and fungi in the soil, as well as roots or other plants that also emit volatile semiochemicals. The complex ecosystem in the soil influences the transport of intraspecific or interspecific information over a greater distance by changing or rather masking the individual volatile blend (Peñuelas et al. 2014). In future, several possible interfering semiochemicals should be tested at the same time with a four-way olfactometer arena. For this, autoclaved millet and the *Metarhizium* spp. isolates should be analysed for the exact composition of the volatile blends including the relative amounts of individual substances by GCMS analysis. Accurate information on the volatile semiochemicals involved in the interaction between wireworms and *Metarhizium* spp. can improve the control strategy by selecting highly virulent isolates with attractive blends. In addition, the AgriMet-Granule could be optimised. A special coating associated with volatile semiochemicals could enhance the attractiveness to ensure contact even under influence of other volatile stimuli.

In conclusion, the contact between the AgriMet-Granule and wireworms is assured even when there is no urge to feed at high conidia concentrations. Millet represents an attractive bait for the AgriMet-Granule formulation of *M. brunneum* intended for wireworm control. However, wireworms’ preference is influenced by the respective *Metarhizum* spp. isolate. In addition, factors influencing the application environment were not considered during the standardised laboratory tests. It is unclear, to what extent the attracting effect of the AgriMet-Granule is still present among numerous other stimuli in the soil. Greenhouse and field application could provide further information to assess the full control potential of the AgriMet-Granule.

## Notes

### Competing Interest Statement

The authors have declared no competing interest.

